# Deep learning-based identification and quantification of rare circulating hybrid cells in orthotopic pancreatic cancer models

**DOI:** 10.64898/2026.08.14.744773

**Authors:** Cody C. Rounds, Divya Ravi, Ge Huang, Biruk Mengesha, Sophia Tran, Allyson Garcia, Nicole Rueb, Young Hwan Chang, Byung S. Park, Melissa H. Wong, Summer L. Gibbs

## Abstract

**Significance:** Rare-cell identification in fluorescence microscopy remains challenging because targets are sparse and background varies between specimens. Combining specimen-specific fluorescence enrichment with image classification may enable efficient and more specific automated detection of rare cells.

**Aim:** We developed a two-stage framework to identify and quantify candidate rare circulating hybrid neoplastic cells (CHCs, ECAD+/CD45+) in peripheral blood mononuclear cell (PBMC) preparations from tumor-bearing and tumor-naïve mice.

**Approach:** PBMCs from 28 mice were imaged by multichannel fluorescence microscopy. Matched unstained samples established animal-specific ECAD and CD45 background distributions for candidate cell enrichment. Blinded multi-annotator consensus labels were used to train a convolutional neural network (CNN) from DAPI, ECAD, and CD45 image crops. Generalization was evaluated by leave-one-animal-out validation across 10 random initializations. Final classification used a 10-model ensemble, and rare-cell burden was compared between groups using negative-binomial regression with total segmented-cell count as an exposure.

**Results:** Of the 1,065,512 segmented cells, enrichment retained 10,176 candidates (0.96%), reducing the search space by >99%. Four of five evaluable tumor-bearing animals showed reproducible held-out discrimination, with median quantified area under the receiver operator characteristic curve (AUROCs) of 0.918–0.951; one animal was a reproducible outlier (median AUROC, 0.338). Ensemble deployment identified 157.94 positive-consensus cells per 50,000 segmented cells in tumor-bearing animals versus 49.55 in controls. The estimated rare-cell rate was 3.15-fold higher in tumor-bearing animals (95% CI, 0.91–10.99; two-sided p=0.071; prespecified one-sided p=0.036).

**Conclusions:** Specimen-specific fluorescence enrichment combined with supervised image classification reduced the cellular search space and enabled automated quantification of a rare CHC (ECAD+/CD45+) phenotypes. Cross-animal validation also identified specimen-specific generalization failure, highlighting the importance of biological-specimen-level validation.

## 1 Introduction

Fluorescence microscopy is widely used to identify and quantify rare cell populations defined by biomarker co-expression, but accurate detection of rare events remains a persistent challenge for the imaging modality.^1–3^ This difficulty is driven in part by target prevalence; rare cells can represent <1% of an imaged specimen. Additionally, autofluorescence and nonspecific staining vary from specimen to specimen, and thus a single fluorescence intensity threshold applied uniformly across specimens often over estimates background in some samples while missing true biomarker-specific signal in other samples.^2–5^ This problem is particularly acute in the detection of circulating hybrid cells (CHCs), a cell population formed by fusion between neoplastic and immune cells that retain functional and phenotypic attributes of both parent lineages and is shed into peripheral blood.^6,7^ As a result, CHCs can be identified by co-expression of an epithelial marker, such as E-cadherin (ECAD) or cytokeratin, with the pan-leukocyte marker CD45.^8^ CHCs were first identified in the peripheral blood of patients with pancreatic ductal adenocarcinoma (PDAC) and have since been detected across multiple cancer types.^6,9–13^ CHC abundance has been shown to be associated with metastatic burden and treatment response in PDAC patients, motivating our interest in establishing an automated imaging pipeline capable of identifying these rare cell types reliably across specimens.^6,9^

We previously developed an image-analysis pipeline for CHC identification in immunofluorescence images of patients’ peripheral blood mononuclear cells (PBMCs), combining a β-variational autoencoder (β-VAE) for single-cell representation learning with a support vector machine (SVM) classifier. That work achieved near-human-level classification accuracy and illustrated the value of specimen-resolved validation.^14^ Unsupervised representation learning via a β-VAE is well suited in settings where the population of interest is not fully specified in advance to allow candidate phenotypes to be discovered from the structure of the data itself rather than defined *a priori*.^15,16^ In the current work the phenotype of interest was known and localized to specific protein markers and fluorescence channels rather than requiring discovery. This allowed us to train a convolutional neural network (CNN) directly on labeled examples of the target phenotype, which prompted use of a supervised CNN in place of the VAE and SVM approach used previously.

We applied this classifier to an orthotopic mouse model of PDAC, which offered experimental control of PBMC collection from tumor-bearing animals that is not available in patient cohorts, making this system well-suited to isolating the effects of orthotopically grown PDAC tumors on CHC burden. CHCs have previously been enumerated in mouse models by manual review of stained PBMC immunofluorescence images, but this approach is time-consuming and does not scale readily to the animal numbers needed for rigorously powered comparisons.^13^ Building on our prior pipeline, we therefore sought to extend automated CHC detection to a mouse model of PDAC. Extending this approach to a new setting also enabled comparison of trained CNN performance consistency across animal PBMC samples rather than only on average since a model can achieve a strong pooled accuracy score while performing poorly on one or more individual specimens. We addressed this inconsistency by validating the classifier on a per-animal basis rather than relying on a single aggregate performance metric.

Herein, we describe a two-stage framework for identifying and quantifying rare CHC candidate cells in multi-channel fluorescence images of PBMCs from an orthotopic PDAC mouse model. Candidate enrichment did not rely on a fixed global intensity threshold or a size-based pre-sort. Instead, each animal’s matched unstained control PBMCs defined its own background distribution in the ECAD and CD45 channels, and candidates were enriched against their animal-specific baseline. These enriched candidates were classified by a CNN trained on blinded, multi-annotator consensus labels. We assessed generalization with leave-one-animal-out validation across ten independently initialized models, and final classification calls were based on agreement across the resulting ensemble. Here, we report the extent to which this enrichment strategy reduced the cellular search space, the consistency of classifier generalization across held-out animals, and whether the resulting ensemble-based rare-cell burden differed between tumor-bearing and tumor-naïve animals.

## 2 Methods

### 2.1 Cell culture & tumor implantation

KPC8060 cells, a murine PDAC cell line, were gifted from Dr. Michael Anthony Hollingsworth (University of Nebraska Medical Center, Omaha, NE). Cells were cultured in Dulbecco’s Modified Eagle Medium (DMEM) supplemented with 10% fetal bovine serum (FBS) and 1% penicillin-streptomycin. At approximately 80% confluence, cells were detached using 0.25% trypsin-EDTA and pelleted by centrifugation at 200 × g. The supernatant was removed, and cells were resuspended at a concentration of 5 × 10³ cells per 50 µL in chilled 1:1 phosphate-buffered saline (PBS):Matrigel (Corning, Corning, NY, USA). The resulting cell suspension was loaded into pre-chilled 1 mL syringes fitted with chilled 28-gauge hypodermic needles and maintained on ice until tumor implantation. Tumor implantation was performed over two consecutive days, with 10 animals undergoing surgery per day. For each surgical session, 600 µL of cell suspension was prepared, corresponding to the volume required for 10 injections plus approximately 20% excess.

All animal procedures were performed under a protocol approved by the OHSU Institutional Animal Care and Use Committee (IACUC; protocol TR04_IP0000202). Ten male and 10 female C57BL/6 mice, 8–10 weeks of age, were purchased from Charles River Laboratories (Wilmington, MA, USA) and allowed to acclimate for 2 weeks in the OHSU vivarium prior to tumor implantation. Animals were therefore approximately 10–12 weeks old at the time of surgery.

Mice were anesthetized by intraperitoneal administration of ketamine: xylazine (100:20 mg/kg), and an appropriate plane of anesthesia was confirmed by toe-pinch response. A left flank incision was made and the peritoneal cavity entered. The spleen was used as an anatomical landmark to identify and isolate the pancreatic tail, which was gently exteriorized with forceps and placed on sterile gauze draped across the lower abdomen. The chilled 28-gauge needle containing the cell suspension was inserted into the pancreatic tail, and 50 µL containing 5 × 10³ cells was slowly injected. Successful intrapancreatic injection was confirmed by formation of a small, localized bleb within the pancreatic tissue.

Following implantation, the pancreas was returned to the abdominal cavity. The peritoneum was closed using absorbable sutures, and the skin was closed with surgical staples. Veterinary-grade cyanoacrylate-based tissue adhesive was applied over the incision to create a watertight seal. Postoperative analgesia was provided by intraperitoneal administration of extended-release buprenorphine emulsion.

Tumor growth was monitored by abdominal palpation. Forty-one days after tumor implantation one mouse was found deceased during routine monitoring. To minimize the risk of additional mortality, the remaining animals were euthanized on the same day by CO₂ inhalation. Euthanasia was immediately followed by collection of approximately 600–800 µL of blood by cardiac puncture, stored in a heparinized vacutainer, followed by resection of the primary pancreatic tumor. Across the cohort, primary tumors had a mean mass of approximately 0.7 g at endpoint. One additional mouse yielded <100 µL of blood during cardiac puncture, which was insufficient for feasible PBMC isolation and was therefore excluded from downstream blood-based analyses. The final cohort available for PBMC analysis consisted of 18 tumor-bearing mice.

### 2.2 PBMC isolation & antibody staining

Prior to animal euthanasia and blood collection, Superfrost Plus microscope slides (Fisher Scientific, Waltham, MA, USA) were fitted with 8-well microarray gaskets and clips (Grace BioLabs, Bend, OR, USA). Two wells were used for each PBMC specimen: one for antibody staining and one as an antibody-negative control. These are referred to hereafter as the “stained” and “unstained” wells, respectively. Slides were coated with 57 µg/mL poly-D-lysine (MilliporeSigma, Burlington, MA, USA) in 1× PBS adjusted to pH 7.4 and incubated for at least 30 minutes prior to PBMC seeding.

For PBMC isolation, whole blood was diluted with 1× PBS to a total volume of 30 mL in a 50-mL conical tube. Thirteen milliliters of Ficoll-Paque Plus (GE Healthcare, Chicago, IL, USA) was then slowly underlaid beneath the diluted blood to establish a distinct interface between the Ficoll and blood layers. Samples were centrifuged at 650 × g for 10 minutes at 4 °C with the centrifuge brake disabled. The PBMC-containing interface was collected using a serological pipette and transferred to a fresh tube, then diluted to a total volume of 40 mL with 1× PBS. Samples were centrifuged again at 650 × g for 10 minutes at 4 °C, with the brake enabled at its lowest setting following centrifugation. The supernatant was carefully aspirated without disturbing the cell pellet, and the cells were resuspended in fluorescence-activated cell sorting (FACS) buffer consisting of 2% fetal bovine serum (FBS), 1 mM EDTA, and 1× PBS adjusted to pH 7.4. Cell concentration and viability were assessed by Trypan blue exclusion using a standard hemocytometer, after which the suspension was adjusted to 50,000 viable cells per 300 µL.

Following poly-D-lysine incubation, the coating solution was aspirated and 300 µL of the PBMC suspension was added to each stained and unstained well. Slides were incubated at 37 °C for 15 minutes to promote cellular adhesion to the poly-D-lysine-coated surface. The cell suspension was then gently aspirated and replaced with 200 µL of 4% paraformaldehyde (PFA) for 5 minutes at room temperature. PFA was removed, and each well was washed once with 200 µL of 1× PBS. The PBS was then aspirated, and the microarray gaskets and clips were removed.

Slides were subsequently permeabilized for 12 minutes in ice-cold cytoskeleton (CSK) buffer containing 100 mM sodium chloride (NaCl), 300 mM sucrose, 3 mM magnesium chloride (MgCl₂), 10 mM piperazine-N,N′-bis(2-ethanesulfonic acid) (PIPES), and 0.5% Triton X-100. Following permeabilization, slides were post-fixed in ice-cold 4% PFA in 1× PBS for 10 minutes and then incubated in ice-cold 2× saline-sodium citrate (SSC) buffer for 3 minutes. Slides were dehydrated through a graded ethanol series consisting of 70%, 95%, and 100% ethanol for 3 minutes each. Prior to antibody staining, the adherent PBMCs were rehydrated by sequential incubation in 100%, 95%, and 70% ethanol, followed by transfer into 1× PBS.

After rehydration, a hydrophobic barrier was drawn around each well to maintain separation between the stained and unstained conditions. Once the barrier dried, both wells were incubated with 2% bovine serum albumin (BSA) in 1× PBS for 30 minutes at 20 °C. Blocking buffer was then aspirated from the stained well, which was incubated overnight at 4 °C with directly conjugated antibodies diluted in the same blocking buffer: FITC-conjugated anti-mouse CD45 antibody (clone 30-F11; BD Biosciences, Franklin Lakes, NJ, USA) at 1:100 and Alexa Fluor 647-conjugated anti-E-cadherin antibody (clone 24E10; Cell Signaling Technology, Danvers, MA, USA) at 1:200. The unstained well was incubated in blocking buffer alone and otherwise underwent identical downstream processing.

Following approximately 16 hours of incubation, antibody and blocking-buffer solutions were aspirated, and slides were immediately washed sequentially in three separate 250-mL baths of 1× PBS for 5 minutes each. Both stained and unstained wells were then incubated with 0.5 µg/mL DAPI in 1× PBS for 10 minutes at room temperature, followed by three additional 5-minute washes in separate 250-mL baths of 1× PBS. After the final wash, slides were promptly mounted using 100 µL Fluoromount-G mounting medium (SouthernBiotech, Birmingham, AL, USA) and 22 × 40 mm #1.5 coverslips (Fisher Scientific, Waltham, MA, USA). Mounted slides were allowed to dry for 24 hours at room temperature under protection from light prior to immunofluorescence imaging.

### 2.3 Immunofluorescence microscopy

Whole-well fluorescence images of stained and unstained PBMC specimens were acquired as 16-bit CZI mosaics at a spatial resolution of 0.325 µm/pixel. Imaging was performed at the OHSU Advanced Light Microscopy Core (ALMC) using a Zeiss Axioscan 7 slide scanner (Carl Zeiss Microscopy GmbH, Oberkochen, Germany) equipped with LED illumination and a Hamamatsu ORCA-Flash sCMOS camera (Hamamatsu Photonics, Hamamatsu, Japan). Images were acquired using a 20×/0.8 NA air objective in the DAPI, FITC/AF488, and AF647 fluorescence channels.

The DAPI channel used excitation and emission bandwidths of 370–410 nm and 430–470 nm, respectively. AF488 fluorescence was acquired using the FITC filter set, with excitation and emission bandwidths of 450–490 nm and 500–550 nm, respectively. The AF647 channel used excitation and emission bandwidths of 625–655 nm and 665–715 nm, respectively.

Autofocusing was performed using nuclear fluorescence in the DAPI channel through sequential coarse-and fine-focus acquisition. Coarse focusing was performed using a 5×/0.1 NA objective at five distinct locations across each slide, followed by fine focusing with the same 20×/0.8 NA objective used for image acquisition. Focus selection was performed using the instrument’s maximum total variation (MTV) autofocus algorithm.

### 2.4 Image processing & cell enrichment

To analyze PBMC fluorescence images at the single-cell level, nuclear segmentation was performed using Mesmer^17^ to generate labeled nuclear masks for each sample. Mesmer was selected over a conventional watershed-based segmentation approach because PBMC density varied substantially among specimens, and densely packed or touching nuclei were incompletely separated by the classical watershed implementation evaluated during pipeline development. The CNN-based Mesmer segmentation provided more reliable nuclear separation under these conditions and was therefore used for the final single-cell analysis.

Segmentation was conducted on both unstained and stained wells. Sparse-field DAPI quality-control safeguards were applied to suppress spurious nuclear detections in low-cell-density regions, and segmented objects lacking sufficient underlying raw DAPI support were removed, where “sufficient” underlying DAPI signal was defined as any nuclear mean intensity below the 5^th^ percentile of nuclear DAPI intensities. A cell-boundary mask was subsequently generated by expanding each accepted nuclear label by one pixel; thus, membrane-marker fluorescence was not used to define PBMC boundaries. The process of enrichment, annotation, training and deployment used in this study is illustrated in Fig. 1. All computational analyses were conducted using custom Python scripts.

**Figure 1.**
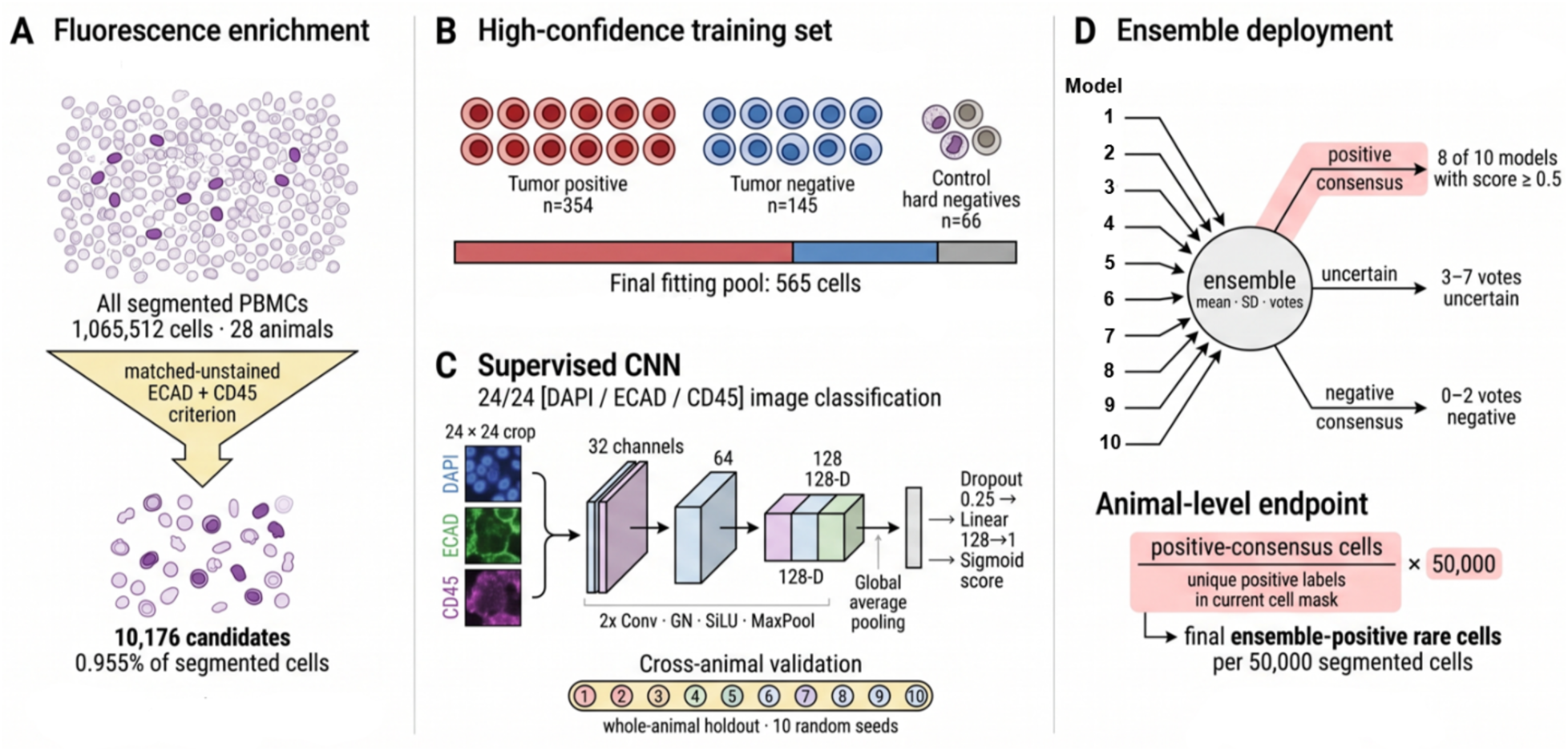
Computational workflow for CHC identification and classification. A) A total of 1,065,512 segmented cells from tumor-bearing animals were screened for candidate CD45+/ECAD+ cells by comparing stained-cell mean fluorescence intensities with animal-specific unstained intensity distributions. B) A subset of enriched candidates, including highconfidence double-positive and definitive negative cells, was manually annotated for CNN training and validation. C) Ten CNN models were trained using whole-animal holdout validation, with each model initialized from a unique random seed. D) Unannotated enriched cells were independently evaluated by all 10 trained CNNs. Cells receiving positive classifications from 8– 10 models were designated positive consensus, those receiving 3–7 positive votes were designated uncertain, and those receiving 0–2 positive votes were designated negative consensus. Only positive-consensus cells were included in final per-animal CHC counts. Figure was created using Biorender.

The stained and matched unstained wells were not registered, and no cell-to-cell correspondence between the two preparations was assumed. Instead, the matched unstained well served as an animal-specific reference for the distribution of background/autofluorescence signal measured in the ECAD and CD45 imaging channels. These distributions were used to define Stage-1 candidate-enrichment thresholds in the corresponding stained well rather than to establish biological marker positivity directly.

For each marker *m*, the nonnegative mean cell intensity *I_i,m_* was calculated for every segmented cell *i* in the matched unstained well and transformed to log space according to

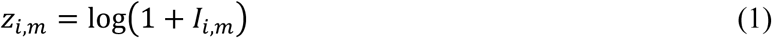

A robust estimate of the spread of the log-transformed unstained distribution *s̃_m_* was then calculated from the median absolute deviation (MAD):

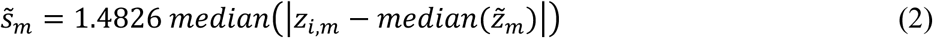

The factor 1.4826 scales the MAD such that it is comparable to the standard deviation for an approximately Gaussian distribution, while retaining the greater resistance of median-based statistics to extreme values.^18^ To prevent nearly invariant unstained distributions from producing spuriously small scale estimates, *s̃_m_* was constrained to be no smaller than the 25th percentile of the positive animal-level scaled-MAD values observed across the cohort for the corresponding marker. The resulting scale estimate was denoted *S_m_*.

The animal-and marker-specific enrichment threshold was then defined at three robust scale units above the median of the log-transformed unstained distribution and transformed back to the original fluorescence-intensity scale:

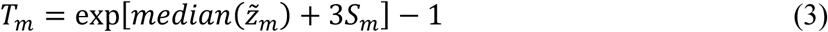

The enriched cell population was then defined as those cells with mean intensities greater than the per-animal threshold set based on the above calculations, leading to a total enriched cell population of 10,176 cells for classification by the CNN.

### 2.5 Human annotation

To facilitate standardized human review of Stage-1 enriched candidate cells, a 36 × 36-pixel image crop was generated for each selected cell and centered on its nuclear centroid. Nuclear centroids were calculated from the corresponding labeled nuclear segmentation masks using custom Python code and SciPy. In addition to the fluorescence channels, a segmentation-derived channel identifying the boundary of the specific cell under review was appended to each image stack. Each candidate was subsequently exported as an individual multichannel TIFF file for annotation. The 36 × 36-pixel crops used for human review were generated specifically to provide annotators with sufficient cellular and local spatial context and were distinct from the 24 × 24-pixel crops subsequently used as inputs to the CNN.

Human annotation were performed by individual annotators who had previously been trained in fluorescence-based cellular phenotyping. This training included recognition of biologically appropriate subcellular localization of marker signal and differentiation of true cellular fluorescence from nonspecific signal, background, overlap from adjacent cells, and imaging artifacts. For the present study, annotators were specifically instructed to determine whether the cell under review demonstrated colocalized, cell-associated ECAD and CD45 fluorescence with signal localized appropriately to the cell membrane. Representative strict positive and strict negative annotations are shown in Fig. 2A and Fig 2B, respectively. Cells lacking convincing membrane-associated signal for either marker, obvious non-cellular artifacts, or in which apparent colocalization could not confidently be attributed to the segmented cell under review, were classified as negative.

**Figure 2.**
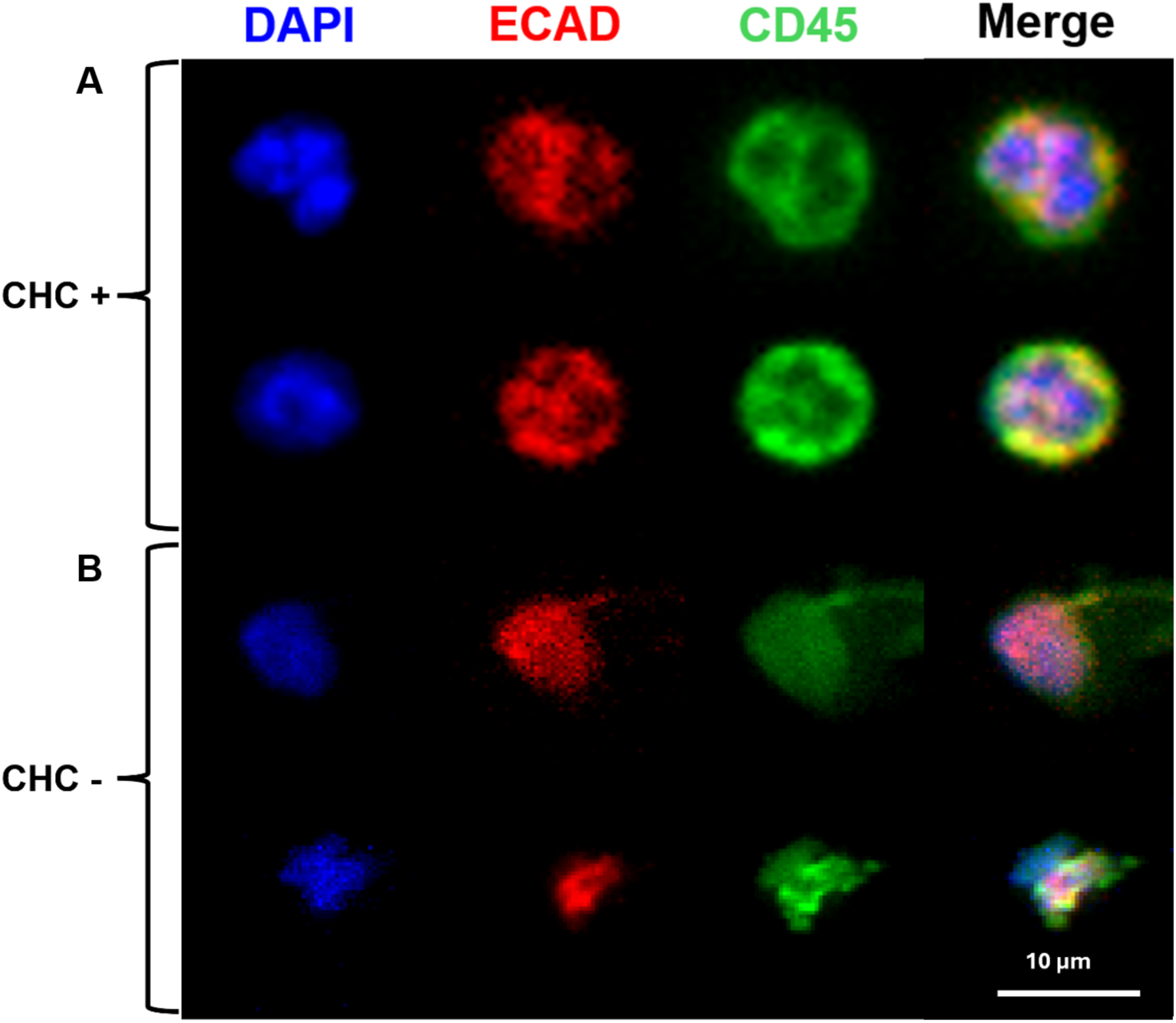
Representative annotated cells used for CNN training. A) Representative cells annotated as CHC-positive. B) Representative cells annotated as CHC-negative. For each example, images are shown from left to right as merged fluorescence, DAPI, ECAD, and CD45 channels.

A total of 700 cells were randomly selected from the Stage-1 enriched candidate population across 5 randomly selected animals for manual review. Each cell was independently evaluated by three to four annotators. Candidate-cell images were imported into a QuPath^19^ project for visualization, and annotation decisions were recorded in cross-referenced Microsoft Excel spreadsheets using the unique cell identifier assigned during segmentation. Annotators were blinded to the originating animal and experimental group, such that cells from tumor-bearing and tumor-naïve animals were evaluated without knowledge of disease status. Annotators evaluated each cell independently and were not provided with the annotations assigned by other reviewers.

Only cells for which all available annotators reached unanimous agreement were assigned a strict positive or strict negative label for supervised model development. Cells with any disagreement among annotators were excluded from the strict training set rather than adjudicated into either class. This approach intentionally prioritized label confidence over training-set size. Of the 700 candidate cells subjected to manual review, 572 satisfied the unanimous-consensus criterion and constituted the high-confidence annotation set used for subsequent classifier development.

### 2.6 Supervised convolutional neural network

Using the open-source Pytorch^20^ package, a directly supervised convolutional neural network (CNN) was trained from three-channel 24 x 24 pixel crops containing DAPI, ECAD, and CD45. No segmentation-mask channel was provided to the final model. The network contained three convolutional blocks with 32, 64, and 128 feature channels, respectively. Each block contained two 3 x 3 convolutions, each followed by GroupNorm^21^ and SiLU activation,^22^ followed by 2 x 2 max pooling. Adaptive global average pooling reduced the final feature tensor to a 128-dimensional image representation. Dropout^23^ with probability 0.25 applied before a linear 128-to-1 output layer.

The network was optimized with binary cross-entropy with logits using AdamW^24^ with a learning rate of 1 x 10^-3^, weight decay of 1 x 10^-4^, and minibatch size 64. Each model was trained for a fixed 20 epochs. Sigmoid transformation of the output logit produced a 0-to-1 classifier score for inference. These scores were treated as classifier evidence and were not interpreted as calibrated biological probabilities.

### 2.7 Fold-specific normalization, augmentation & class/animal balancing

Image-intensity normalization was calculated independently within each training fold to prevent information leakage from the held-out animal.^25^ For each training animal and channel, the 99.8^th^-percentile of target-cell pixel intensity was calculated. The median of these animal-level percentiles across training animals defined the fold-specific scale for that channel. Each pixel value was divided by the corresponding training-derived scale and clipped to the interval [0,1]. Held-out animals were transformed using only normalization parameters calculated from the training animals.

Training augmentation consisted of random rotations by multiples of 90 degrees, random horizontal and vertical reflection, independent multiplicative channel-intensity jitter from 0.90 to 1.10, and occasional low-amplitude Gaussian noise. Augmented inputs were clipped to [0,1].

To reduce domination by heavily annotated animals or by the majority class, samples were drawn with combined animal-and class-aware weighting. Each cell was initially weighted approximately inversely to the number of labeled cells contributed by its animal, using an animal-count floor of five cells. Class-level scaling was then applied so that positive and negative examples contributed approximately equal sampling mass. Sampling with replacement was used during each training epoch.

The complete experiment was repeated across 10 independently initialized random seeds. Random seeds were treated as technical optimization replicates rather than independent biological replicates. The primary validation quantity was the distribution of held-out-animal receiver operator characteristic (ROC) curve area under the curve (AUC)^26^ values across seeds and animals rather than a pooled ROC curve formed from outputs of independently trained fold models.

### 2.8 Classification metrics

For each held-out animal, threshold-independent discrimination was summarized by area under the receiver operating characteristic curve (AUROC). Average precision (AP)^27^ was used to characterize retrieval of positive cells. Because the no-skill AP baseline is dependent on positive-class prevalence, the baseline was defined as the proportion of positive cells within each held-out animal. AP lift was calculated as AP minus positive prevalence, such that positive AP lift indicated positive-cell ranking above the specimen-specific no-skill baseline.

Binary classification performance was additionally evaluated using a prespecified sigmoid-score threshold of 0.5. Sensitivity, specificity, precision, F1 score, balanced accuracy, and accuracy were calculated independently for each evaluable held-out animal. Sensitivity and specificity were considered together to determine whether the fixed threshold consistently balanced detection of positive and negative cells across animals or instead favored one class in particular specimens. This analysis was used to assess the consistency of the fixed decision threshold across specimens and was considered complementary to the threshold-independent AUROC and AP analyses.

### 2.9 Final multiseed ensemble deployment & visualization

The final deployment ensemble consisted of the exact 10 validated final models produced during the multiseed experiment. Each of the 9,476 unannotated Stage-1 candidate cells were independently scored by all 10 CNNs. Per-cell outputs included all individual model scores plus the ensemble mean, median, standard deviation, minimum, maximum, range, and vote concordance.

An individual model cast a positive vote when its sigmoid score was at least 0.5. Operational ensemble categories were defined as positive consensus for 8-10 positive votes, uncertain for 3-7 positive votes, and negative consensus for 0-2 positive votes. These categories were used as reproducible deployment rules and not as biologically calibrated probability thresholds. The ensemble mean sigmoid score was likewise interpreted as a model score rather than a calibrated probability of cellular identity.

For visualization of the learned image representation, the 128-dimensional CNN representation after adaptive global average pooling was projected into two dimensions with UMAP.^28^ The same coordinates were displayed according to ensemble score, animal identity, training role, and ensemble consensus. Representative microscopy examples were selected algorithmically from the learned representation rather than by manual visual selection. UMAP was used as a qualitative visualization; absolute two-dimensional (2D) distances were not interpreted as quantitative biological distances.

### 2.10 Animal-level rare-cell burden

The final biological endpoint was calculated at the animal level from positive-consensus ensemble calls. For animal i, rare-cell burden, B_i_, was defined as the number of positive-consensus candidates, Y_i_, divided by the total number of segmented cells, N_i_, in the current same-animal cell mask and multiplied by 50,000 to yield approximate counts per 50,000 segmented cells.

The denominator, *N_i_,* was defined as the number of unique cell labels in the segmentation mask for that animal. Candidate counts, annotation counts, feature-table row counts, and the maximum numerical cell label were not used as substitutes for total segmented-cell count because label identifiers contained gaps due to the initial DAPI-based hallucination filtration. The animal, rather than the individual cell, was the biological unit for downstream group inference.

### 2.12 Statistical analysis

The final tumor-versus-control analysis modeled the raw positive-consensus cell count using a negative-binomial NB2 regression^29^ with the logarithm of each animal’s total segmented-cell count as an exposure offset. The model was defined as:

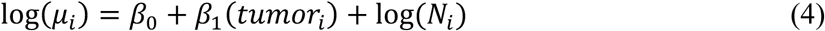

with NB2 variance as:

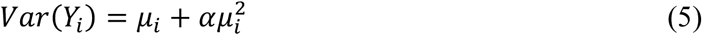

The exponentiated tumor coefficient was interpreted as the tumor-to-control incidence rate ratio (IRR) for ensemble-positive rare cells.

A Poisson regression model with the same exposure offset was fit as a diagnostic for overdispersion. Poisson adequacy was evaluated using the Pearson dispersion statistic, with values substantially greater than 1 indicating variance exceeding that permitted under the Poisson assumption. Because substantial overdispersion was observed, the negative-binomial NB2 model was used for primary inference. Uncertainty in the NB2 IRR was additionally assessed with 5,000 animal-level bootstrap^30^ in which whole animals were resampled within biological groups. Sensitivity analyses of normalized per-50,000 burden included Welch’s unequal-variance t-test, Mann-Whitney U testing, and a 100,000-replicate animal-label permutation test^31^ of the difference in group means.

Given the original hypothesis that tumor-bearing animals have a higher prevalence of CHCs (i.e., ECAD^+^, CD45^+^ double-positive cells) relative to tumor-naïve animals, both conventional two-sided tests and prespecified one-sided tests for tumor greater than control were reported. The one-sided permutation probability was calculated directly from the upper tail of the permutation distribution rather than by halving the two-sided probability. The magnitude and uncertainty of the tumor-versus-control difference were emphasized over whether any individual test crossed a p = 0.05 threshold. Welch’s t-test, Mann–Whitney testing, and permutation testing were used as complementary sensitivity analyses because they evaluate different aspects of the group distributions.

## 3 Results

### 3.1 Matched-unstained cell enrichment reduces the PBMC search space

The two-stage analysis combined quantitative fluorescence enrichment with supervised image classification. Across the 28-animal analysis cohort, 1,065,512 segmented cells were screened. Application of the final animal-specific matched-unstained ECAD/CD45 enrichment rule retained 10,176 Stage-1 candidates, corresponding to approximately 0.96% of the segmented-cell population and reducing the search space by > 99% before CNN classification.

### 3.2 Cross-animal validation reveals strong performance in four animals and a reproducible outlier

Four of the five held-out animals showed strong and reproducible ranking performance across independently initialized models as demonstrated by ROC curves across the independently initialized CNNs (Fig. 3A). Median hard-negative AUROC across the 10 seeds was 0.945 for animal 2, 0.946 for animal 3, 0.951 for animal 4, and 0.918 for animal 5. AUROC was at least 0.80 in 10/10 seeds for animal 2, 9/10 for animal 3, 10/10 for animal 4, and 10/10 for animal 5 (Fig. 3B).

**Figure 3.**
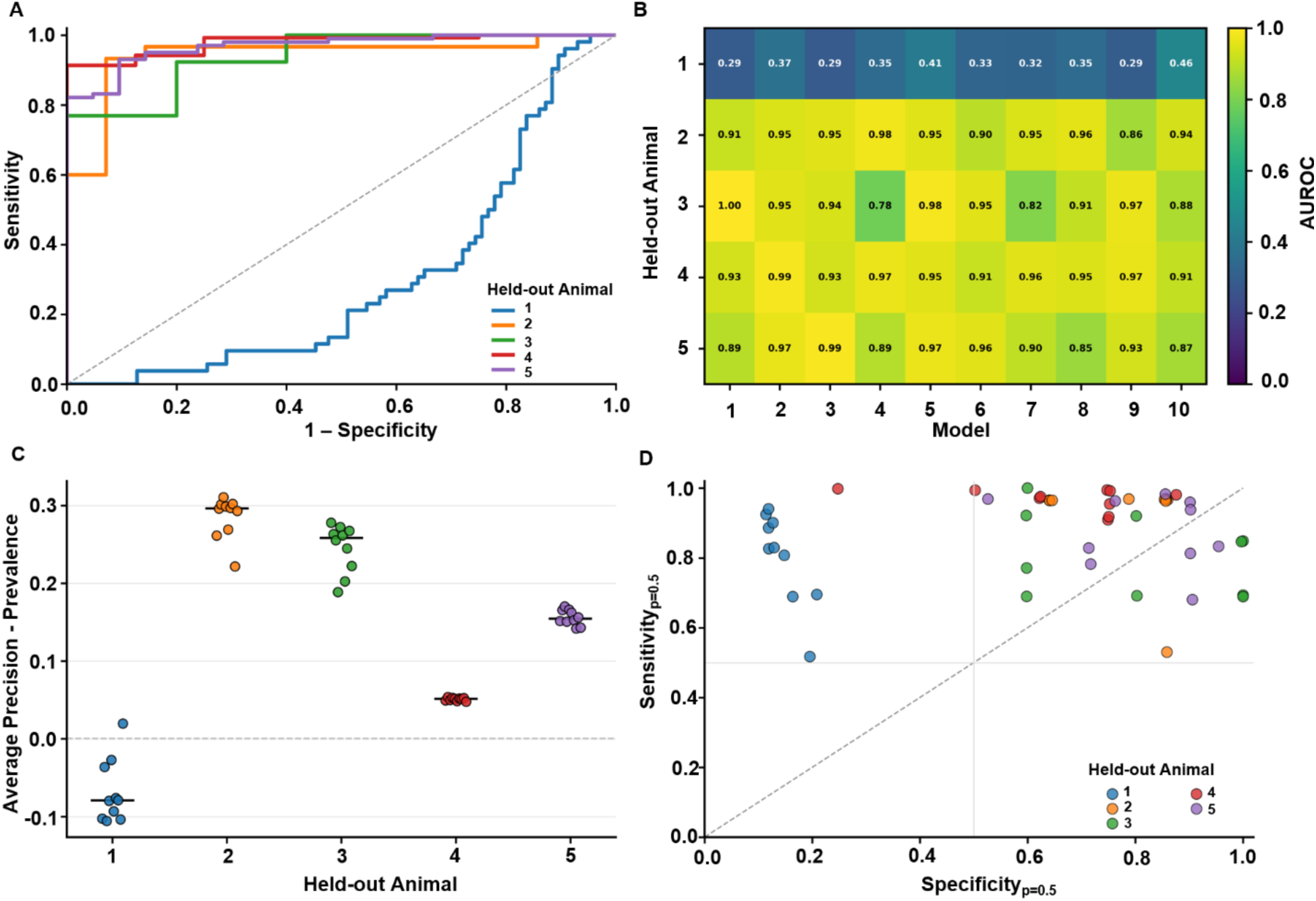
Final CNN ensemble validation and operating-point performance. A) Receiver operating characteristic (ROC) curves for each held-out animal across the 10-model ensemble. B) Corresponding area under the receiver operating characteristic curve (AUROC) values for each held-out animal across the 10 models. C) Average-precision lift above class prevalence for each held-out animal across the 10-model ensemble. D) Sensitivity and specificity analysis for each held-out animal across the 10-model ensemble using a classification threshold of 0.5.

Animal 1 was a reproducible outlier. Its median hard-negative AUROC was 0.338, and none of the 10 seeds achieved AUROC of at least 0.80 (Fig. 3B). Because the failure was reproduced across independent initializations, animal 1 was retained and flagged as animal-specific performance outlier rather than excluded as an unfavorable random seed, allowing the model to learn from the annotations.

Precision-recall analysis was interpreted relative to each held-out animal’s positive prevalence and can be seen for each animal across all model initializations (Fig. 3C). This was important because class composition varied substantially among evaluable animals. In animal 1, AP lift above prevalence was near zero or negative in multiple seeds, whereas the better-generalizing animals demonstrated positive AP lift. Operating-point analysis at a score of 0.5 further showed that ranking performance and binary decision behavior were not interchangeable; some held-out specimens displayed a strong sensitivity-specificity imbalance consistent with specimen-specific score shifts (Fig. 3D). However, across all held-out animal and model initialization combinations, the mean F1 score at this threshold was 0.83, which was aligned with our previously established model-based performance and similar to human-level CHC calling accuracy.^14^

### 3.3 Multi-CNN ensemble deployment provides an auditable measure of model agreement

The final 10-model hard-negative ensemble scored 9,476 Stage-1 candidates. Individual model scores were retained for every cell together with ensemble mean, median, variability and model agreement. Requiring at least 8 of 10 models to cross the 0.5 operating threshold created a conservative positive-consensus category, while cells receiving 3-7 positive votes were retained as uncertain rather than forced into a definitive biological class.

This ensemble formulation separated classifier evidence from model stability. Cells with similar mean scores could be distinguished according to whether independent initializations agreed or produced substantial score dispersion. The resulting consensus categories therefore provided an auditable operational deployment rule while avoiding the stronger claim that sigmoid values represented true biological probabilities.

### 3.4 The learned CNN representation organizes candidate cells by classifier behavior

To examine how Stage-1 candidate cells were organized within the learned CNN feature space, the 128-dimensional image representations were projected into two dimensions using UMAP (Fig. 4). Classifier scores varied systematically across the learned representation (Fig. 4A), where regions of cell classifications with high model agreement there was consistent grouping between tumor-bearing and tumor-naïve mice (Fig. 4B). The locations of the strict positive and negative human annotations demonstrated how the labeled training examples sampled this feature space (Fig. 4C). Mapping the final ensemble classifications onto the same coordinates showed the distribution of positive-consensus, uncertain, and negative-consensus cells across the learned representation (Fig. 4D). To relate regions of the learned representation to the underlying fluorescence images, representative candidate-cell crops were mapped back to their locations within the score-colored UMAP (Fig. 5). Examples were selected to span distinct classifier behaviors, including ambiguous cells, difficult positive and negative examples, and strongly negative cells. The learned representation contains visually interpretable examples spanning model certainty and phenotype difficulty.

**Figure 4.**
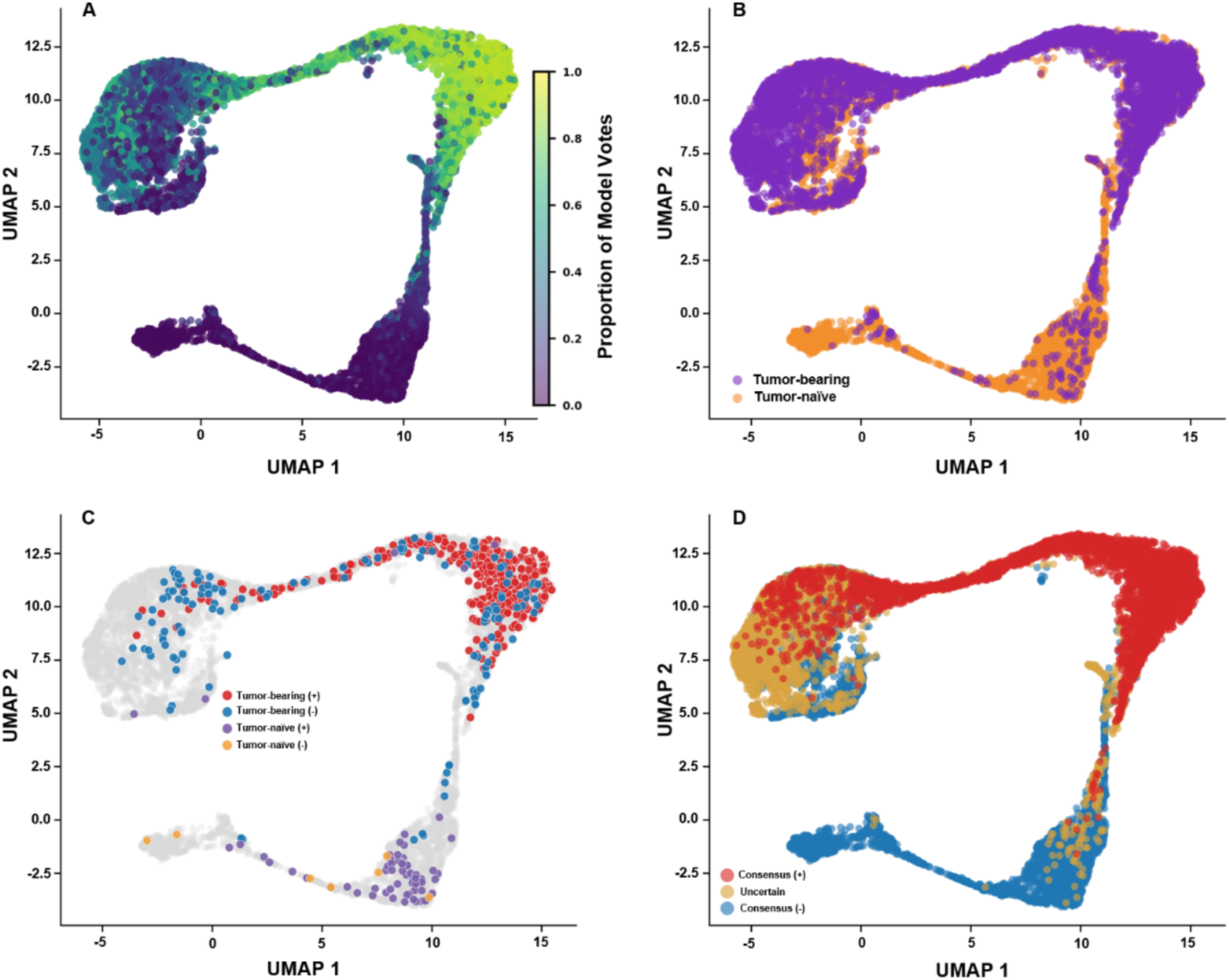
UMAP visualization of CNN embedding and ensemble classifications. A) Representative wo-dimensional UMAP projection of the 128-dimensional CNN embedding for all enriched cells, colored by the number of positive votes received from the 10-model ensemble. B) Distribution of cells from both tumor-bearing and tumor-naïve throughout the embedding space. C) Localization of manually annotated cells within the embedding. D) Final ensemble classification of all enriched cells across the cohort as positive consensus, uncertain, or negative consensus.

**Figure 5.**
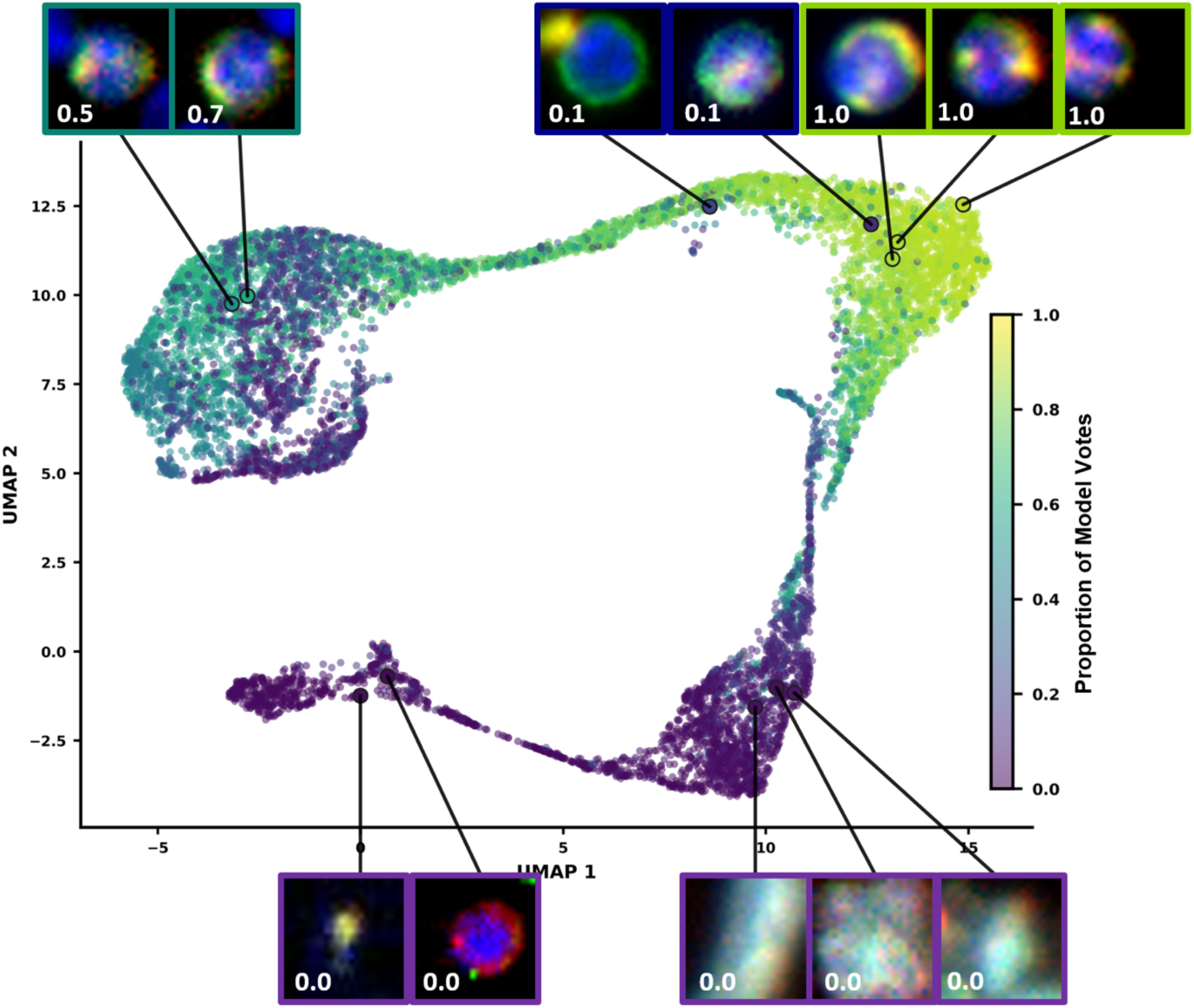
Representative cell morphologies across CNN embedding space. Representative cells sampled from locations across the CNN-derived UMAP embedding. In the top row, the first two cells were classified as uncertain, receiving 5 and 7 positive votes out of 10, respectively; the third represents a negative-consensus cell; and the final two represent positive-consensus cells. The bottom row contains representative negative-consensus cells sampled from additional regions of the embedding. UMAP is colored according to the number of positive ensemble votes received by each cell, as in Figure 4A.

### 3.5 Tumor-bearing animals exhibit an increased estimated burden of CHCs

Animal-level rare-cell burden was summarized as the number of positive-consensus cells per 50,000 segmented cells (i.e., CHCs). Tumor-bearing animals had a mean burden of 157.94 positive-consensus cells per 50,000 segmented cells compared with 49.55 per 50,000 in controls animals, while median burdens were 126.26 and 21.63 per 50,000, respectively (Fig. 6)

**Figure 6.**
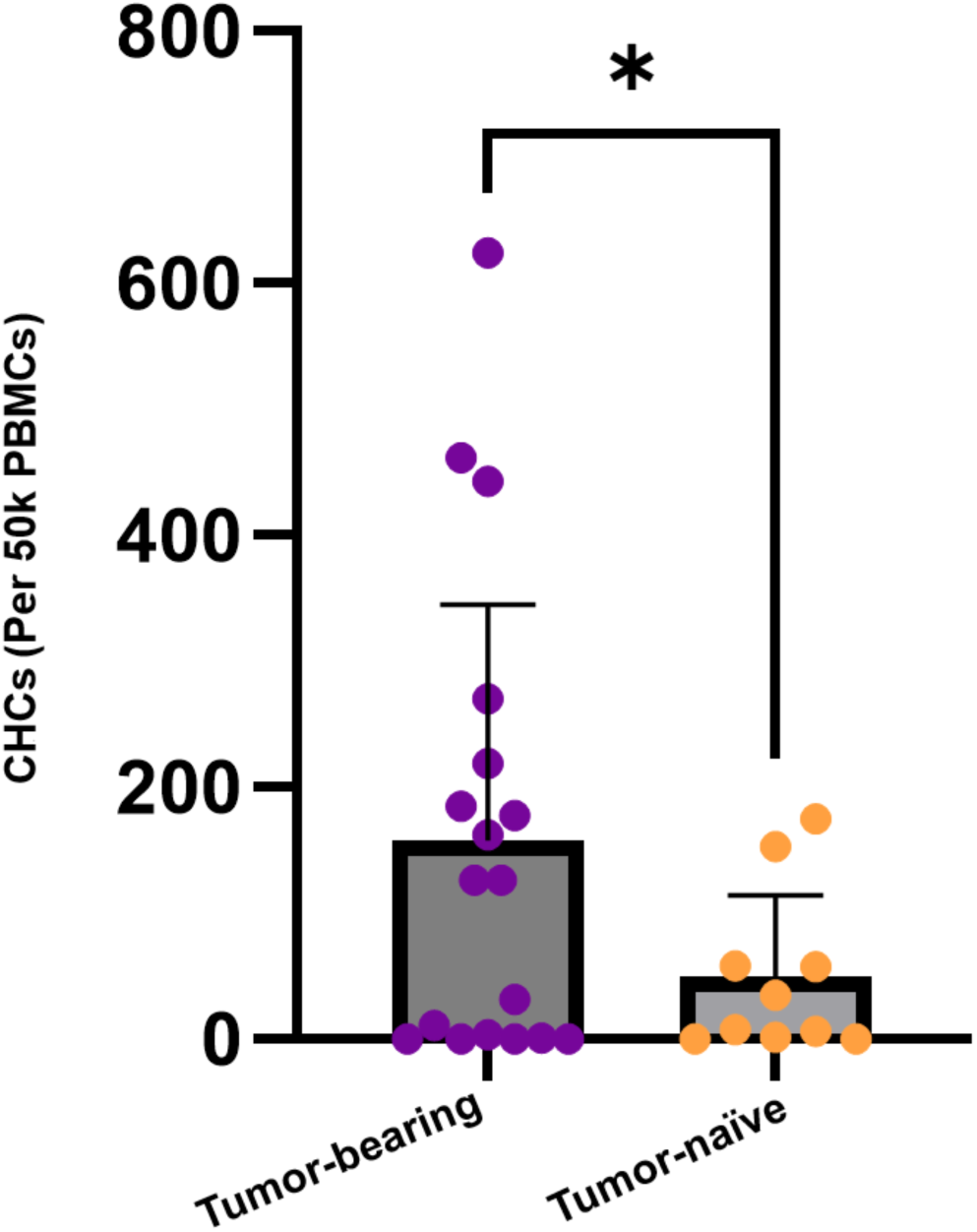
Final CHC counts in tumor-bearing and tumor-naïve animals. CHC counts are compared between tumor-bearing [Tumor (+)] and tumor-naïve [Tumor (-)] animals. From left to right, data are shown as violin plots depicting the distribution of per-animal counts, individual-animal values illustrating cohort-level spread, and group means with ±1 standard deviation. All y-axes report CHC counts per 50,000 cells. Significance denoted: p = 0.036, defined via the one-sided NB2 regression analysis.

For primary statistical inference, raw positive-consensus cell counts were modeled using negative-binomial NB2 regression, with each animal’s total number of segmented cells included as an exposure offset. The model estimated a 3.15-fold higher rate of ensemble-positive cells in tumor-bearing animals relative to controls (IRR = 3.15; two-sided 95% CI, 0.91-10.99; p = 0.071). Because the prespecified directional hypothesis was that tumor-bearing animals would have a higher rare-cell rate than controls, the corresponding one-sided test was also evaluated and yielded p = 0.036, with a one-sided 95% lower confidence bound for the IRR of 1.11.

Bootstrap resampling of whole animals produced a two-sided 95% confidence interval for the IRR of 1.32-10.76. The count data showed marked overdispersion relative to a Poisson model, with a Pearson dispersion statistic of 81.7. Consistent with this finding, the NB2 model fit the data substantially better than the corresponding Poisson model (Akaike Information Criterion = 264.8 versus 2532.4).

Secondary animal-level tests gave mixed results. Welch’s unequal-variance t-test gave two-sided p = 0.036 and prespecified one-sided p = 0.018. The animal-label permutation test gave two-sided p = 0.086 and one-sided p = 0.034. Mann-Whitney testing gave two-sided p = 0.258 and one-sided p = 0.129. Together, these analyses indicate a higher estimated rare-cell rate in tumor-bearing animals, while also demonstrating substantial animal-to-animal variability and overlap between the two groups.

## 4 Discussion

Using animal-specific enrichment followed by ensemble CNN classification, we investigated performance for identifying and quantifying rare CHCs in an orthotopic PDAC mouse model. Enrichment against each animal’s own background distribution reduced the candidate pool from greater than 1×10^6^ million segmented cells to roughly ten thousand candidates, making downstream classification manageable at the scale needed for a large, multi-sample comparison. Cross-sample validation evaluated each held-out animal individually rather than relying on a single pooled performance metric, revealing a reproducible failure across all ten models in a single animal. The result points to a broader issue for rare-cell classifiers. Aggregate validation metrics can hide meaningful differences in model performance across individual specimens and reporting performance only in aggregate risks overstating classifier accuracy.

The classifier’s constrained model likely contributed to its consistent performance across animals. Since the phenotype of CHCs are well established, a directly supervised CNN could be trained on a well-defined classification problem rather than on an open-ended representation learning task. The smaller more constrained CNN model had fewer parameters to overfit to any single specimen’s heterogeneity, which may also explain why four of five animals generalized well despite a comparatively small and consensus filtered training set.

Animal-specific enrichment additionally contributed to model performance. A single global intensity threshold would have forced a tradeoff. Set conservatively, it would risk discarding true candidates in cleaner specimens. Set permissively, however, would risk flooding the candidate pool with background-driven false positives in nosier ones. Defining each animal’s threshold relative to its own matched unstained well avoided this tradeoff by letting the enrichment criterion adapt to each specimen’s own noise floor rather than forcing every animal through the same autofluorescence cutoff. This strategy addresses variation between animals but did not address variation within a single stained sample. Autofluorescence and nonspecific staining are not necessarily uniform across a stained sample. For example, a candidate cell located in a brighter region of the image for reasons unrelated to true marker expression could pass the enrichment threshold. Correcting for within-image variability would require co-registered, pixel-wise background subtraction.

Performance was strong for four of the five animals in the training set, with AUROC values consistently >0.80 across all ten models. The fifth animal, in contrast, showed AUROC values near or <0.34 across every model. The consistency of this failure across ten independent models argues against random noise but rather suggests a systematic difference the model could not accommodate. Staining quality can vary for multiple reasons during sample preparation, which could shift the appearance of both positive and negative cells in ways the model had not encountered during training on the other animals. Unusually high or unevenly distributed background in this particular specimen could have enriched atypical candidates before reaching the classifier. This result importantly illustrates why validation at the level of individual animals is necessary. A pooled AUROC would have neglected to inform the complete failure on the fifth animal since it would reflect an average dominated by the majority. Leave-one-animal-out validation repeated across multiple random initiations is what made this failure visible. We report it here to make the case that specimen-level validation should be standard for classifiers applied to small and heterogenous cohorts.

We also report a small number of CHCs in tumor-naïve animals, which raises the possibility that these cells appeared in healthy controls. As previously mentioned, co-registered background subtraction was not performed in this study, which would have resulted in a fraction of calls reflecting retained autofluorescence rather than true signal. In addition, low levels of CHCs have also been reported in healthy patient controls.^9^ Whether the double positives observed here reflect this same baseline population, residual background, or both cannot be resolved using the current dataset. Distinguishing between these possibilities would require co-registered background subtraction or a CHC-specific marker capable of separating genuine hybrid cells from background artifact.

The 3.15-fold increase in CHCs called in tumor-bearing mice is consistent with prior findings in human PDAC samples, where CHC abundance has similarly been associated with presence and burden of disease.^6,9^ The statistical support for this difference was mixed rather than uniform. The negative binomial rate ratio reached significance under a one-sided test, but other statistical tests did not agree. This likely reflects the same animal-to-animal variability discussed previously rather than an absence of a true effect since one animal was found to be unreliable at the classification stage. A small sample size with a single outlier animal can compound this effect. Nonetheless, these results support increased CHC counts in tumor-bearing mice, which are consistent with what is found across multiple cancer subtypes in patients.

Several limitations should be considered regarding the classifier’s training data and specificity. First, the marker panel used for CHC classification was limited to ECAD and CD45. Although the two-marker definition is consistent with prior CHC literature, it cannot fully distinguish genuine tumor-immune fusion events from other sources of double positivity, including macrophages that have phagocytosed epithelial or tumor material without undergoing fusion.^32–34^ Additionally, the classifier was trained on consensus-labeled cells from a subset of the full cohort. Requiring unanimous agreement between annotators improved label confidence but further reduced the number of available training examples. A larger sampled training set would help clarify whether the generalization failure observed in one animal reflects a genuine specimen-level difference or a limitation of training data coverage.

Taken together, these results demonstrate that specimen-level validation and animal-specific enrichment are both necessary and achievable for rare-cell quantification in preclinical PDAC models. Animal-specific enrichment made automated CHC quantification achievable in a mouse model without relying on presorting for target cells or a fixed global threshold. Beyond CHCs, our framework offers a template for other rare-event classification challenges in fluorescence imaging. In the context of PDAC, our work also establishes CHC burden as a reliable and automated readout in future studies of tumor-immune interactions in PDAC.

## Acknowledgements

We would like to thank the Advanced Light Microscopy Core and the Brenden-Colson Center for Pancreatic Care at Oregon Health and Science University. We also acknowledge Anna Igler for technical assistance with antibody validation, PBMC slide preparation, and sample annotation. We further thank Brandon Pham and Nathan Mebratu for annotating samples. Figures were created using Biorender.

